# Growth and transgenerational acclimatization of juvenile *Pocillopora damicornis*

**DOI:** 10.1101/2020.11.02.364596

**Authors:** Lev Gerstle

## Abstract

Global carbon emissions and associated increase in ocean temperatures are understood to be the main driving force in the degradation of coral reefs. Elevated temperatures impact various life stages of scleractinian corals, from the free-floating planulae of brooding corals to older, sexually viable individuals. With global warming, questions have arisen over whether organismal adaptation will be enough to keep up with the pace of environmental change. Researchers have pursued investigations of whether or not rapid acclimatization, through transgenerational plasticity, can help protect populations until genetic adaptation occurs. Acclimatization in corals has been widely studied in all life stages of corals, with the important exception of recently settled juveniles. In this study, I built upon past research by exposing adult *Pocillopora damicornis* colonies to elevated (28.5°C) or ambient (25.5°C) temperatures and examining the settlement ability and growth of their planulae *ex situ*. Juveniles from preconditioned parents fared better in higher temperatures compared to their naïve counterparts. Lunar timing of planula release between treatments peaked at different times in the lunar cycle. Peak planula release occurred on lunar day 23 for prestressed corals and on lunar day 7 for corals from ambient temperature seawaters. While future projects should follow up on these preliminary trials with *in situ* experiments to assess this phenomenon in the field, this study represents an important step in understanding how corals may be able to acclimatize and eventually adapt to climate change.

## Introduction

Scleractinian corals are vulnerable to changes in seawater temperatures caused by global climate change. Living just 1°C–2°C below their upper thermal limit, corals are responsive to even slight shifts in temperatures (1). Average global ocean temperatures are predicted to increase 2°C–3°C by the end of the century, meaning that declines in reef health due to annual bleaching events are unlikely to abate in the next few decades (2). These well-documented biological responses highlight the need to understand how corals intrinsically react to environmental shifts (3,4). A broad avenue of research has opened up focusing on the potential for trans-generational acclimatization in some coral species.

All organisms experience at least some level of environmental heterogeneity; sessile species, unable to move to areas with more favorable conditions, need to develop mechanisms to increase odds of survival. One such mechanism is plasticity, which can occur between generations as parental conditions influence offspring characteristics (5). Some of the earliest and best studied examples of plasticity comes from plants, with both genetic and phenotypic plasticity cited as common maternal influences (6). With future climate scenarios predicted to impact all life stages of corals, it is important to understand how cross-generational plasticity might improve the responses to some of these stressors at individual points in the growth process. Chua *et al.* (2013) found that temperature influenced early life history stages of corals, while Albright and Langdon (2011) showed the same life stages were influenced by both temperature and ocean acidification (OA). Foster *et al.* (2016) found that elevated PCO_2_ reduced skeletal weight of coral recruits and caused the formation of deformed and porous skeletons. Post-recruitment growth and skeletal formation are critical to maintaining healthy reefs, and each of these studies in isolation shows a potential point of disruption in the life cycle of corals.

The question then becomes to what extent acclimatization can accommodate any one of these stressors in isolation, or potentially multiple points of disruption simultaneously. Investigations into transgenerational plasticity in other marine invertebrates has yielded mixed results. Calcifying tubeworms exposed to different pH levels had varying degrees of acclimatization based on gender, with egg laying females having less energy stored away for response to acidification (10). *Tripneustus gratilla*, the common collector urchin, has shown the ability to acclimatize to both acidification and ocean warming based on parental environment. However, this ability comes with the tradeoff of a reduction in both larval arm length and egg size from individuals raised in heated seawaters (11). These results point to tradeoffs in a fast shifting environment.

Corals have been found to be able to acclimatize to temperatures via preconditioning within a single generation (12). Remarkably, this can happen within a single yearly cycle and has been documented *in situ* on large portions of the Great Barrier Reef. In 2016, exposure to heat levels capable of causing stress elicited a 50% probability of severe bleaching, while in 2017 the same 50% response occurred at much higher heat levels over a longer period of time. Responses in 2017 were also contingent on the degree of thermal stress that corals were exposed to in 2016 (13). While these single generation studies are illuminating and important, they fail to capture the full complexity of coral systems. Genetic and epigenetic feedback loops have the ability to generate phenotypic plasticity, and both can alter an organisms response to environmental change (14). This proves an especially difficult puzzle to solve with corals due to both their longevity and the length of time required to reach reproductive fertility. Few studies have been conducted at a cross-generational scale, although the last few years have produced several of note. High temperatures combined with OA treatments negatively affected adult corals, and caused individuals to produce larvae with variable sizes and metabolic conditioning when subsequently re-exposed (15). *In situ* testing found that brooded larval survivorship was 14% higher in corals growing in elevated temperatures on Lihuitou Reef in China with *p*CO_2_ and temperature synergistically enhancing carbonic anhydrase activity, a key enzyme involved in photosynthesis (16).

To date, much of the limited research that has been conducted at a cross-generational scale has focused on the larval stage. A missing element is a focus on the post-settlement juvenile stage. Environmental changes that impact this stage have potential long-term implications for skeletal growth (9). Additionally, research that addresses how parental pre-conditioning effects multiple life stages is sparse. In this study, I tested whether preconditioning of brooded planulae resulted in any measurable trans-generational acclimatization in the newly released larvae. I compared larvae held at 25.5°C (hereafter known as ambient) to larvae held at 28.6°C (hereafter known as elevated). I also tested whether bleaching history changes timing and amplitude of planula release. I used *Pocillopora damicornis* as a study species, which provides an ideal test system as they brood their offspring internally (15), release planulae year round (17), and have a fixed lunar release cycle (16).

## Methods

### Collection

*Pocillopora damicornis* were provided by the State of Hawaii’s Division of Aquatic Resources Hawai’i Coral Restoration Nursery. Corals were originally collected from an area between Pier 41 and Sand Island Bridge in Honolulu Harbor. Collection occurred on four separate dates between February and April of 2019. Twenty adults were randomly chosen from the collection and divided into two subgroups.

### Experimental Design

On May 17^th^, 2019, corals were placed into two 150-liter indoor tanks within a closed system and with artificial lighting. Tanks were equipped with sand, carbon, and UV filters as well as a sediment trap and multiple bag filters. Lighting was set to ~100 μmol m^−2^ s^−1^, the recommended acclimation setting at the Coral Restoration Nursery. Power heads were installed in each tank and set to 50% constant flow to aid in water circulation. Two 1000-watt titanium heaters with electronic temperature gauges were used to heat the tanks. Temperatures were set to 25.5°C for both tanks and corals were allowed to acclimate for one week. Water flow for each tank was kept between 4-6 l min^−1^. Temperature in one tank was then raised by 1°C every week for three weeks, until the water temperature reached 28.5°C. This temperature was then maintained for three weeks, until signs of stress began to appear. Stress indicators included prolonged polyp withdrawal, minor sloughing of tissue, and bleaching. Heaters were then turned off and temperatures were ramped down to 25.5°C over a weeklong period. E_PAR_ levels, temperature, and flow were measured daily with a PAR meter, electronic and mercury thermometers, and measurement of 6 seconds of water flow into a beaker, respectively. Corals were fed a combination of two brine shrimp cubes, a single nutrient puck comprised of *Spirulina*, mysids, and *Artemia*, as well as ~175 ml of Continuum Aquatics Coral Exponential Amino Acids (Continuum Aquatics, Fort Payne, AL, USA) every Monday, Wednesday, and Friday.

### Planula Collection

Corals were transferred to an outdoor planulation table on June 17^th^, 2019. Each coral was placed into an individual plastic mixing bowl, so that flowing water allowed the buoyant planulae to flow over the smooth handle and into collection beakers with plankton mesh sides. Beakers were checked every morning for planulae, and planula release was tracked for 3.5 months. Planulae were transferred into and stored in 1000ml Erlenmeyer flasks and kept from settling with magnetic stir plate and stir bar. Seawater in the flasks was changed every week. Water temperature on the planula table was measured daily and fluctuated between ~26 and 27°C. Mean lunar day of release of planulae was determined using circular statistics. Statistical analysis was performed in R(18).

### Settlement

Four 5-liter tanks were used for settlement of planulae. Seawater from a supply well was filtered using a bag and sand filter in series. Aragonite tiles, which had been cured in flowing sea water for the three months, were arranged in a 3 by 4 grid on the bottom of the tanks. The bottoms of the tanks between the tiles were filled with sand to encourage settlement on tiles. Holes were drilled into the side of the tanks and covered with plankton mesh to allow water to flow through the tanks. Water flow was kept at ~1.5 l min^−1^. Planulae were divided into one of four groups: planulae from preconditioned parents to be settled in heated seawater (n=7), planulae from preconditioned parents to be settled in ambient temperature seawater (n=8), planulae from naïve parents in heated seawater (n=46), and planulae from naïve parents in ambient temperature seawater (n=46). Two header systems were built to allow even heating of seawater using two 1000-watt titanium heater. On settlement day, seawater in the tanks was lowered to one-third capacity and flow was turned off to ensure planulae did not settle on the upper glass of the tanks and were not washed through the tanks. Temperatures were taken every fifteen minutes with an electronic thermometer. Water level was kept at one-third capacity from 11am to 4pm, at which point flow was restored.

### Juvenile Measurements

After one month of growth, juvenile corals were photographed weekly using a Moticam X^3^ Wi-Fi camera (Motic, Kowloon Bay, Kowloon, Hong Kong) and compound microscope. Growth was subsequently measured for 6 weeks using built in Moticam software. After six weeks, photophysiology of Symbiodinacea was measured as the maximum relative rate of electron transport (ETR) for a standardized 40 μmol m^−2^s^−1^. Photosynthetic efficiency was measured during rapid light curves (RLC) using a pulse-amplitude-modulated (PAM) fluorometer (Diving-PAM, Walz GmbH, Germany; methods drawn from (19) and (20)) and WinControl-3.30 software with saturating pulses at 0:10 intervals and the following increasing actinic light values: 10, 20, 30, 40, 50, 60, 70, 80, 90, 100 μmol photons m^−2^ s^−1^. The intensity of the red light-emitting diode (655 nm peak, <0.15 m^−2^s^−1^, at 1.6 kHz) was set to 8 and was too low to induce fluorescence when used as a probe light. Maximum fluorescence (F_m_) was measured during a saturating light pulse from a halogen lamp (0.8 seconds, at >5,000 μmol m^−2^s^−1^; KL1500, Schott, Mainz, Germany) exposed to the sample via an optical fiber. Aragonite tiles were scrubbed free of algal growth around the new recruits to prevent measurement errors. The diving-PAM probe was fitted with a black plastic tube to minimize light leakage. Corals were re-measured using the diving-PAM 48 and 96 hours after initial measurements. Missing ETR values, likely a result of low baseline values (F_0_ or F_t_), were manually calculated using the equation:

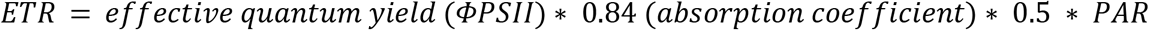

where Φ_PSII_ = (F_m’_–F_t_)/F_m’_ or Y(II). Relative ETR (rETR), calculated as:

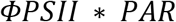

was used in addition to PAM ETR output values because the absorption coefficient for corals is likely different than that of plants (21,22). Juvenile growth and health were analyzed using a two-way ANOVA with fixed factors of parental preconditioning status and juvenile environment and their interaction. Statistical analysis and data transformation were carried out using Python (23).

## Results

Peak planula production for prestressed adults occurred on lunar day 23 with a circular standard deviation of 8.4, while peak production for naïve adults occurred on lunar day 7 with a circular standard deviation of 3.6 (Fig 1). A two-way ANOVA test determined that pre-conditioning of parental corals was found to be independently significant examining its effect on juvenile growth (p-value = 0.04; Table 1), while temperature of current environment (ambient or elevated) was not a significant factor (p-value = 0.20; Table 1). However, it was found that there was no significant interaction between environmental temperature and preconditioning on juvenile growth (p-value = 0.80; Table 1). Change in area was, on average, higher in juveniles from preconditioned adults for both the elevated and ambient temperature groups (Fig 2). Meanwhile, growth at ambient temperature was higher than growth in elevated temperatures for both the preconditioned and naïve juveniles (Fig 2).

**Table 1.** The effects of stress on coral growth.

| Source of Variation | df | F | PR(>F) |
| --- | --- | --- | --- |
| Change in Area (mm<sup>2</sup>) |  |  |  |
| Adult Preconditioning History | 1 | 5.101 | **0.0474** |
| Experimental Temperature | 1 | 1.870 | 0.2013 |
| Preconditioning*Temperature | 1 | 0.063 | 0.8066 |
| ETR |  |  |  |
| Adult Preconditioning History | 1 | 10.538 | **0.0070** |
| Experimental Temperature | 1 | 20.739 | **0.0006** |
| Preconditioning*Temperature | 1 | 1.932 | 0.1897 |
| rETR |  |  |  |
| Adult Preconditioning History | 1 | 5.782 | **0.033** |
| Experimental Temperature | 1 | 7.653 | **0.017** |
| Preconditioning*Temperature | 1 | 0.040 | 0.843 |

The effects of the interaction of adult preconditioning history and experimental heat stress on change in area, electron transport rate (ETR), and relative electron transport rate (rETR). Juvenile response analyzed with two-way ANOVA. Bold indicates significance.

**Figure 1.** Planula Release. Timing of planula release per lunar days in June -October 2019, from both naïve and preconditioned temperature parental corals.

**Figure 2.** Response of juvenile P. damicornis to heat stress following pre-conditioning. Larvae were pooled by preconditioning treatments and held at either elevated or ambient temperature prior to settlement. Juveniles were tracked for 8 weeks before measuring: a) growth rate as a change in area; b) electron transport rate using PAM fluorometry; c) relative electron transport rate calculated using PAM fluorometry. Significant effects of preconditioning were found for ETR, growth rate and rETR. Growth rate, ETR, and rETR were lower after elevated temperature exposure for both groups, but higher on average in the preconditioned group than the naïve group.

For ETR values, both preconditioning of parental corals (p-value = 0.007; Table 1) and temperature of current environment (p-value = 0.0006; Table 1) were found to be independently significant of each other. There was no significant interaction between environmental temperature and preconditioning on ETR (p-value = 0.189; Table 1). As with growth rate, ETR was higher in juveniles from preconditioned adults for both the elevated and ambient temperature groups, and also higher at ambient temperature for both naïve and preconditioned juveniles (Fig 2).

For rETR, both preconditioning of parental corals (p-value = 0.03; Table 1) and temperature of current environment (p-value = 0.017;Table 1) were also found to be independently significant of each other, similar to ETR. There was no significant interaction between environmental temperature and preconditioning on rETR (p-value = 0.843; Table 1). rETR was higher in juveniles from preconditioned adults for both high and ambient temperature groups, and also higher at ambient temperatures for both naïve and preconditioned juveniles (Fig 2). This followed similar patterns to both ETR and growth rates.

## Discussion

Observations of growth in juvenile *Pocillopora damicornis* allowed investigation of stress responses in a critical, yet understudied, stage of coral development. Additionally, measurements of photobiological health in juveniles using PAM fluorometry introduces a potential new technique for the assessment of growth in the early life of corals, as this technology had previously been used mostly for the assessment of adult coral health. Growth was higher in recruits from preconditioned parents, as well as in both preconditioned and naïve juveniles in ambient temperature seawaters (Fig 2). The same trends were seen for rETR, with higher levels in preconditioned individuals as well as in both preconditioned and naïve juveniles grown in ambient seawaters. rETR also had less variability than growth numbers, indicating that this assay may have been a better indicator of relative health of juveniles than a simple measurement of juvenile area. These results provide evidence that bolsters the idea that cross-generational acclimatization is a process that has the potential to mitigate the effects of a changing climate on multiple life stages of corals.

rETR of corals settled in elevated temperature treatments was significantly higher for juveniles from preconditioned parents compared to juveniles from naïve parents. Two potential mechanisms could explain a trans-generational shift that would cause differences between the naïve and preconditioned juveniles. The first is the development rate of brooding planulae and the timing of planula release. Temperature has been shown to reduce the number and timing of reproductive events in *Pocillopora damicornis* (24). The peak days of planulation for prestressed adults occurred on lunar day 23. This average was likely driven by the single large planulation event on the 23^rd^ lunar day in August. Naïve adults, meanwhile, had a peak release on lunar day 7 (Fig 1). They also had lower, more consistent peak release numbers, rarely releasing more than 25 planulae at a time. Reproductive processes in corals have been shown to be plastic (25), and it is possible that this plasticity contributed to acclimatization in juveniles. By the time planulae were released, peaking later in the month for prestressed corals, any traits that would have failed to survive higher temperature conditions would have been weeded out, as planulae possessing these traits failed to survive the brooding process.

A second possibility is that, rather than acclimatization weeding out unfit planulae during brooding, adult corals passed resistance directly to offspring. Temperature stress has been shown to reduce important biochemical components in adult corals such as lipids, proteins, and mycosporine-like amino acids, the reduction of which was amplified in offspring from bleached parents (26). A reduction in lipid content in *P. damicornis* also coincided with a decrease in planulae and gametes during planulation events (27). It is possible that the vertical transmission of Symbiodinacea from parent colonies conditioned against temperature stress was picked up by PAM fluorometry and manifested as the change in rETR values observed here. If thermal stress limited which corals were able to planulate due to a lack of sufficient resources to put toward reproduction, this might provide another explanation for why the planulae that came from pre-stressed corals had higher photochemical responses.

There are other pathways of acclimatization that might explain the difference between the responses of corals from naïve and preconditioned treatments but were outside the scope of this experiment to test. One such direction for future research is the role of mycosporine-like amino acids (MAA). These molecules are small, water soluble, UV radiation absorbers found in a wide variety of marine organisms (28). Historically, research on MAAs has focused on their usefulness as UV protectors (29–31). However, evidence has emerged that they could additionally act as secondary metabolites, serving a multipurpose role (28). A 2009 study by Yakovleva et al. found that some MAAs provide rapid protection during thermal stress before antioxidant enzymes are activated (32). Typically, the costs associated with symbiosis are not examined, as it is thought to be a mutualistic process that benefits both parties. However, this study found that *Acropora intermedia* larvae containing zooxanthellae showed decreases in specific MAAs thought to reduce oxidative stress, decreases in survival rates, and increases in oxidative cellular damage when subjected to elevated temperatures. Because this study was conducted on such an early life stage, there is a compelling case that MAAs might explain at least some of the rapid acclimatization to thermal stress in *Pocillopora damicornis* and other coral species.

It is possible that tank effects could have influenced the outcomes of this study and that the experimental design involved pseudoreplication, since all adult corals were held in just two separate tanks and juveniles were held in just four separate tanks. However, due to limitations in available resources necessary to maintain corals in individual tanks, the relatively small biomass of corals used coupled with large tank volume and high water flow (4-6 l min^−1^ for adult tanks, 1.5 l min^−1^ for juvenile tanks) should have minimized any effects of pseudoreplication. Growth of algae in the juvenile tanks could have also led to an increase in dissolved organic carbon, facilitating microbial growth and reducing oxygen availability for juveniles. However, daily cleaning coupled with high flow and regular turnover of water in the tanks should have been enough to keep microbial buildup below harmful levels.

There were limitations to this investigation that could be improved upon in future studies. Among treatments, numbers of planulae settled were uneven, with two groups of n=46 planulae settled from naïve parents, and two groups of n=7 and n=8 planulae settled from preconditioned parents. Unequal planulation among adult corals is a well understood problem of multigenerational experiments on corals, and one of the unique challenges with investigating transgenerational plasticity across multiple generations and life stages. Additionally, questions arose over the use of PAM fluorometry as an appropriate technique for the measurement of juvenile health. Photosymbiotic algae are integral to reef building corals growing in oligotrophic waters, as they provide photosynthate to help meet daily nutritional demands (21). The development of PAM fluorometry techniques has allowed for a non-invasive measurement of the relative health of corals by quantifying metrics such as the potential quantum yield of photosystem II and the estimated rates of photosynthetic electron transport, but the majority of testing to date has been performed on adult corals. PAM fluorometers measure the relative quantum yield of Chl *a* by targeting samples with excitation pulses, and probing the probability of light energy being re-emitted (19). The ETR_max_ measured here, which peaked somewhere between 40 to 60 μmol m^−2^s^−1^ for all samples, is the maximum amplitude at which a re-emission response was recorded. For measurements taken on December 16^th^, sufficient data were obtained to produce ETR and Y(II) values, which were calculated automatically by the diving-PAM. However, for subsequent measurements taken 48 and 96 hours later, the diving-PAM began to regularly fail to automatically calculate ETR and Y(II). Using the equations Φ_PSII_ = (F_m’_–F_t_)/F_m’_ and rETR = Φ_PSII_ * PAR made it possible to fill in these gaps manually. Values calculated for December 18^th^ and 20^th^ closely matched rETR and ETR outputs from December 16^th^ calculated automatically by the PAM fluorometer, but this most likely oversimplifies relative photobiological health. Fluorescence was significantly lower in these subsequent tests, and because Φ_PSII_ is calculated as a ratio of these two numbers ((F_m’_-F_t_)/F_m’_), this would not have shown up in the Y(II) values. Instead, calculated Y(II) values would have masked any sharp reductions in F_m’_ and F_t_, which would have indicated changes in photobiological processes. Although there is no known evidence to support the potential for damage to juveniles from saturating light pulses (>10,000 μmol photons m^−2^s^−1^ (22)), investigation of this topic could produce interesting and novel results. More work is clearly needed to determine if the efficacy of using PAM fluorometry as a technique extends to coral individuals at the juvenile stage.

As this study drew heavily from elements included in a seminal paper from Putnam and Gates (2015), it is important to elaborate on the similarities and distinctions between the two. The 2015 study examined adult *Pocillopora damicornis* to investigate the effects of heat and ocean acidification on future offspring tolerances. Exposure to elevated temperature and high CO_2_ caused reductions in the photophysiology of adult corals, measured as maximum quantum yield of PSII (dark-adapted F_V_/F_M_) using a PAM fluorometer. Larvae from both high and ambient colonies were then collected, pooled, and reciprocally crossed into either high or ambient environments, mirroring the conditions that stressed the parent colonies. Size differences, dark respiration rates, and size normalized dark respiration rate were then used as metrics of larval health to compare the offspring groups.

This study utilized similar techniques for stress of adult corals, collection of larval offspring, and reciprocal crossing of juveniles into environments mirroring those that stressed adult colonies. What makes this study unique is the examination of possible acclimatization of offspring in the juvenile stage of the coral life cycle and the use of PAM fluorometry as a measurement technique on juveniles. Although it has been documented that *P. damicornis* undergoes sexual spawning (33), examination of populations in Hawai’i indicate the dominance of a small number of clones and a propensity for asexual brooding (34). Offspring from *P. damicornis* undergo exact inheritance of adult genotypes (35), and this brings into question whether variation in offspring sizes and photobiological processes documented here and in the aforementioned Putnam and Gates (2015) paper demonstrates true trans-generational acclimatization. Torda et al. (2017) raises further questions regarding environmental exposure and overlap between generations such as that which occurs in brooding *P. damicornis*, emphasizing the difficulty in teasing apart whether variations in offspring are a result of trans-generational acclimatization or developmental plasticity. Examining trans-generational acclimatization in corals that sexually spawn might solve some of these issues, and while limitations exist in our ability to spawn corals *ex situ* (36), recent breakthroughs in in the propagation of an F2 generation suggests more robust studies of acclimatization across generations are not far off (37). In the meantime, adding tools to our repository of knowledge on brooding corals and acclimatization will help drive knowledge forward as we work to perfect trans-generational study techniques.

Rapid acclimatization in corals provides a bright spot to counter an increasingly bleak view of the future of coral reefs. This study adds to a growing body of evidence that corals are capable of, and possibly succeeding in, acclimatization to various anthropogenic stressors. Further investigation of the wide variety of mechanisms behind this process is critical to the future of coral research. In addition, improvement of techniques to allow multigenerational as well as multi life stage studies of corals is needed to fully understand the long-term effects of acclimatization, and to determine what, if any, negative tradeoffs might occur as a result of the acclimatization process.

## Acknowledgements

I would like to thank the entire team at the State of Hawaii’s Division of Aquatic Resources Hawai’i Coral Restoration Nursery for providing me with the tank space and technical support needed to complete my project: David Gulko, Norton Chan, Chelsea Wolke, Christina Jayne, Callie Stephenson, Raquel Gilliland, Honor Weber, and Taylor Engle. Thank you to my committee for putting up with me and for all of the advice, support, and wisdom you imparted on me during my graduate schooling. A truly special thank you to Dr. Cynthia Hunter, who gave me a lab, a home, and a bit of a push in the right direction. And finally, thank you to Cristina Kennedy, who was my inspiration for going to graduate school in the first place.

## Supporting information

**S1 Fig. Progress of** *Pocillopora damicornis* **juvenile growth.** One representative coral from, top to bottom: preconditioned in ambient temperature seawater, preconditioned in elevated temperature seawater, naïve in elevated temperature seawaters, and naïve in ambient temperature seawaters. Left column is each coral on 11/15/2019, paired with the same coral on 12/16/2019 in the right column. Note differences in scale bars in each image.

